# Revealing abrupt transitions from goal-directed to habitual behavior

**DOI:** 10.1101/2023.07.05.547783

**Authors:** Sharlen Moore, Zyan Wang, Ziyi Zhu, Ruolan Sun, Angel Lee, Adam Charles, Kishore V. Kuchibhotla

## Abstract

A fundamental tenet of animal behavior is that decision-making involves multiple ‘controllers.’ Initially, behavior is goal-directed, driven by desired outcomes, shifting later to habitual control, where cues trigger actions independent of motivational state. Clark Hull’s question from 1943 still resonates today: “Is this transition abrupt, or is it gradual and progressive?”^1^ Despite a century-long belief in gradual transitions, this question remains unanswered^2,3^ as current methods cannot disambiguate goal-directed versus habitual control in real-time. Here, we introduce a novel ‘volitional engagement’ approach, motivating animals by palatability rather than biological need. Offering less palatable water in the home cage^4,5^ reduced motivation to ‘work’ for plain water in an auditory discrimination task when compared to water-restricted animals. Using quantitative behavior and computational modeling^6^, we found that palatability-driven animals learned to discriminate as quickly as water-restricted animals but exhibited state-like fluctuations when responding to the reward-predicting cue—reflecting goal-directed behavior. These fluctuations spontaneously and abruptly ceased after thousands of trials, with animals now always responding to the reward-predicting cue. In line with habitual control, post-transition behavior displayed motor automaticity, decreased error sensitivity (assessed via pupillary responses), and insensitivity to outcome devaluation. Bilateral lesions of the habit-related dorsolateral striatum^7^ blocked transitions to habitual behavior. Thus, ‘volitional engagement’ reveals spontaneous and abrupt transitions from goal-directed to habitual behavior, suggesting the involvement of a higher-level process that arbitrates between the two.

## Main text

Humans and other animals are often thought to be creatures of habit. When driving, for example, we are initially told that the color of a traffic light should guide our actions: green to ‘go’ and red to ‘stop’. Through practice, we learn purposefully and are driven by the conscious goal to follow the rules of the road. Over time, these rules become automatic; without deliberation, we will push the gas pedal on a green light and the brake pedal for a red light. This transition from goal-directed to habitual control has long been assumed to be gradual^8–14^. More specifically, an initial goal-directed action (R) in response to a cue (S) yields a desired outcome (O) which then slowly evolves into a habit where the cue elicits the action (S-R) without necessarily having the goal in mind^15,16^. The automatization of decisions can be thought of as an efficient way to offload well-learned contingencies to free up resources for more flexible, goal-directed learning^17^. The formation and perseverance of habits, however, can also be maladaptive with neural circuits being co-opted in substance use disorders or compulsive behaviors^18–20^. Understanding the exact time course of habit formation is critical to disentangling its neural basis and could help inform future interventional strategies for combating habit-related disorders.

The assumption of a slow, gradual shift from goal-directed to habitual control underpins current models of learning and informs most approaches to understanding the neurobiological basis of habit formation. To date, however, the nature, timing, and properties of the transition between controllers have been challenging to pinpoint due to methodological constraints. In rodents, the gold standard for assessing whether a behavior is under goal-directed or habitual control at a specific time point exploits the observation that goal-directed actions are sensitive to the outcome^21,22^ (e.g., animals will only perform an action when the reward is desired) while habitual behavior is less sensitive to the outcome (e.g., animals will continue to perform said action even if the reward is not explicitly desired). This behavioral characterization relies on defining habitual behavior as the loss or absence of goal-directed control^23,24^. Sensitivity to the outcome has been successfully operationalized in laboratory testing with ‘outcome devaluation’^18,25^ procedures in which a reward is devalued (through satiety or taste aversion). Outcome devaluation, or a related alternative called contingency degradation, is typically implemented at set time points outside of the normal training regimen (e.g., the middle and end of a multi-day training period). To date, no approach exists to disambiguate between goal-directed and habitual control in real-time, during training^26,27^. A complementary approach in the study of habit formation in rodents is to exploit distinct reinforcement schedules to bias goal-directed, or habitual behavior^28^. Animals under distinct reinforcement schedules are then tested for habitual behavior with outcome devaluation. The use of outcome devaluation, contingency degradation, and distinct reinforcement schedules, though powerful, inherently limit assessing the nature, timing, and properties of the transition between goal-directed and habitual control in individual animals due to the discrete test sessions and cohort-level comparisons. Can we behaviorally identify habitual transitions in real-time and during training? Is the transition slow or sudden? What are the characteristics of these transitions in individual animals? Addressing these questions requires a new behavioral approach that assesses the decision mode *en passant*, without discrete ‘test’ sessions, without biasing behavior to one or the other process, and without impacting the ongoing learning process.

Here, we present such an approach. We reasoned that animals are usually highly motivated to perform tasks because water and/or food are restricted and only made available during the task. This leads to a sustained ceiling motivation driven by the animals’ need to obtain their daily food or water intake in a short period of time. In such situations, animals remain highly engaged throughout a task irrespective of the underlying decision mode. We hypothesized that if animals were motivated mainly by a taste preference, rather than a biological need, we could track naturalistic fluctuations in their motivation for the preferred reward by fostering variability in reward-seeking. Under goal-directed control, task engagement would wax and wane naturalistically (‘volitional engagement’), due to palatability-driven motivation, which would lead to reduced responding to the reward-predicting cue; in contrast, under habitual control (in which animals are less sensitive to the outcome), the S-R nature of the behavior would drive high and stable responding to the reward-predicting cue despite ongoing changes in the underlying desire for the outcome.

### A palatability-based approach reduces motivation to consume water

In the home cage, we gave mice *ad libitum* access to water laced with citric acid (CA) (**Fig. 1a_i_**, CA yellow) which makes it slightly acidic to the taste but still fulfills hydration needs^4,5^. Before instrumental training, CA mice lost significantly less weight than mice on a standard water restriction (WR85) protocol (**Fig. 1a_ii_**) (WR85 = 17.8%±2.3%, CA = 8.9%±1.9% average and std weight loss respectively, p=0.000055, Wilcoxon rank sum test). This difference was maintained throughout training (**Extended Data Fig. 1a**) (Wilcoxon rank sum test, p=0.000055), while no differences were observed in the animals’ initial weight (**Extended Data Fig. 1b**) (Wilcoxon rank sum test, p=0.97). Before mice begin discriminative auditory training (**Fig. 1a_i_**), they first experience 2 days of instrumental training in which they learned to make an instrumental response (lick) to subsequently receive a small reward (‘lick training’, 3 µl plain water droplet) (**Fig. 1a_i_**, lavender). During this initial session, we assessed licking patterns as a proxy of motivation. CA mice executed fewer licks per session (**Fig. 1b-c**, and **Extended Data Fig. 1c**, yellow) (Wilcoxon rank sum test, p=0.021), exhibited strikingly different lick patterns (**Fig. 1c**), and obtained significantly fewer rewards compared to WR85 (**Fig. 1d**) (Wilcoxon rank sum test, p=0.00079) while maintaining the same lick frequency when engaged in licking (**Extended Data Fig. 1d**) (Wilcoxon rank sum test, p=0.67). This suggests that under the CA protocol, mice exhibit reduced motivation for plain water.

**Figure 1.**
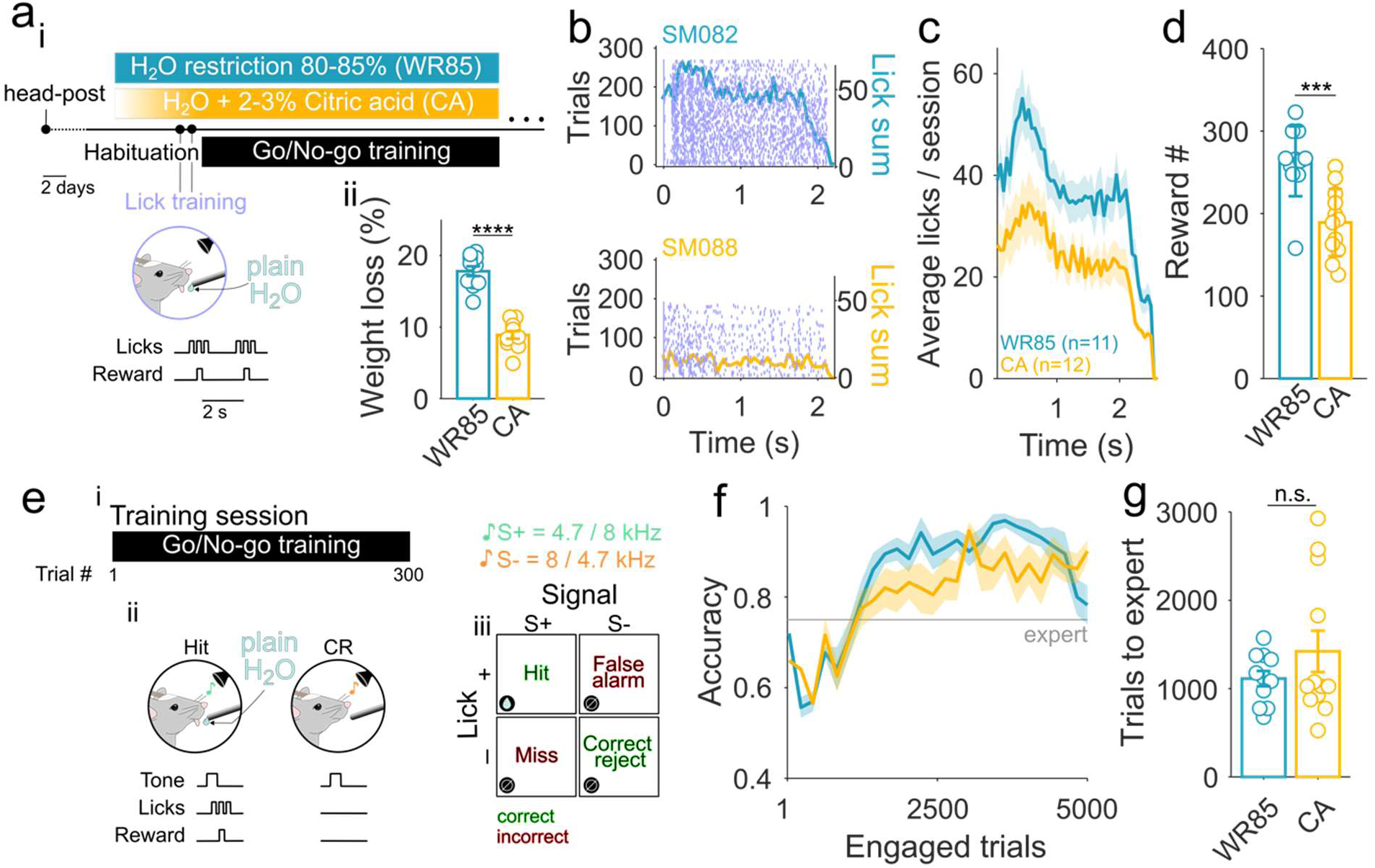
Palatability-based motivation reduces water consumption without impacting learning trajectories. **a_i_,** Protocol outline: after head-post implantation, mice underwent a habituation period and introduction to the water restriction paradigms. ‘Control’ mice underwent a common (maintenance at 80-85% original body weight) water restriction paradigm (WR85, blue). CA mice had *ad libitum* access to a water bottle with a low percentage of a less palatable hydration source (citric acid laced water) in their home cage (CA, yellow) to which they were progressively introduced (starting with 0.5%, reaching maximum 3%). After a few days, both cohorts underwent two days of lick training (lavender), followed by an auditory Go/No-Go training (black). **a_ii_,** CA mice lost significantly less weight compared to WR85 (Wilcoxon rank-sum test, p=0.000055, WR85 n=11, CA n=12). **b,** Lick raster plots during lick training for a WR85 (upper) and a CA (lower) exemplar animal. The right y-axis (color lines) represents a PSTH-like sum of licks for each animal. **c,** Average PSTH-like licking in one session, showing that CA mice lick less (n=11 WR85, and n=12 CA mice). **d,** CA mice obtain significantly fewer rewards compared to WR85 (Wilcoxon rank sum test, p=0.00079) in one lick training session (n=11 WR85, and n=12 CA mice). **e_i_,** After lick-training, mice underwent auditory cued go/no-go training that consisted of ∼300 trials per session. **e_ii_,** Mice learn to lick after a S+ tone to obtain a plain water reward (3ul) and withhold licking to an S-tone to avoid a time-out. **e_iii_)** Correct responses are hits (licking to the S+ tone) and correct rejects (withhold licking to the S-), while incorrect responses are false alarms (licking to the S-) and misses (not licking to the S+). **f,** Accuracy comparison between WR85 (blue) and CA (yellow) mice on highly engaged trial blocks is similar (group comparison ANOVA, F(1,20)=3.06, p=0.081, interaction group x trials F(1,20)=0.83, p=0.68) (n=11 WR85, and n=12 CA mice). Expert accuracy is defined as 75% correct (gray horizontal line). **g,** No differences were observed in the number of trials to reach expert accuracy between groups (Wilcoxon rank sum test, p=0.42) (n=11 WR85, and n=12 CA mice).

### Abrupt transitions from goal-directed to habitual behavior spontaneously occur during training

Mice were then trained on a discriminative auditory go/no-go task in which they learned to lick to one tone (S+, reward-predicting cue) for a water reward (hit) and withhold licking to another tone (S-, non-reward-predicting cue) to avoid a timeout (**Fig. 1e**, correct reject). CA mice exhibited lower response rates to the S+, consistent with the reduced motivation, but surprisingly only minor differences in discrimination performance throughout learning. Specifically, when restricting performance assessment to blocks of high engagement (>50 trials with >90% hit rate), task performance during learning was similar to WR85 mice (**Fig. 1f-g**). In addition, CA and WR85 mice exhibited similar, high discrimination performance (75%) within 1,500 trials (WR85=1132 ± 87 and CA=1422 ± 234 trials, Wilcoxon rank-sum test p=0.93). Overall, these data show that CA mice learn task contingencies at similar rates to WR85 mice while responding less to reward-predicting cues.

We next sought to explore in detail the impact of reduced motivation on responses to the reward-predicting cue. This could be driven by a continuously lower response rate (generally lower motivation) or, alternatively, a fluctuating response rate (periodic changes in motivation). In WR85 mice, animals initially increase their action rates to both tones, followed by a slow reduction in response to the S- (**Fig. 2a**, left example animal). The S+ response (hit rate) stayed consistently high with minimal variability. We observed a striking contrast in CA mice (**Fig. 2a**, right), where for thousands of trials, CA mice exhibit a fluctuating hit rate, regularly shifting from epochs of high hit rate to low hit rate, suggesting that mice are volitionally engaging in the task and that we can track their fluctuating motivation levels. We thus focused on the hit rate as the response rate of interest (**Fig. 2b**). CA mice showed significantly fewer blocks of high engagement (hit rate > 90%) compared to WR85 mice (**Fig. 2c**, Wilcoxon rank sum test, p=0.0028). Surprisingly, after this high-variability phase, most CA mice abruptly transitioned to a low-variability phase (**Fig. 2d_i_**, red line). We observed that 10 out of 12 (83.3%) of the CA mice exhibited high hit rate variability, of which 8 out of 10 (80%) transitioned to a low-variability phase (**Fig. 2e**). Transitions occurred more than 2,000 trials after animals exhibited expert discrimination performance, suggesting this transition was not due to ongoing contingency learning (d’>2 expert=1436±454 trials vs. transition=3832±385 trials) (**Fig. 2f**). To confirm that this change in hit rate was not due to a sudden change in underlying motivation, we measured the weights of the animals daily. Importantly, we observed (1) no differences in discrimination performance around the transition (paired t-test, p=0.83) (**Extended Data Fig. 2a**) (2) no evidence of changes in weight pre- versus post-transition (**Extended Data Fig. 2b**) (paired t-test, p=0.19), and (3) no changes in consumption in the home cage based on measurements of post-task and next-day weights (**Extended Data Fig. 2c**) (paired t-test, p=0.81). This suggests that CA mice do not exhibit increased motivation post-transition and, instead, points to the rapid emergence of habitual control. Interestingly, we observed this transition typically occurred at the beginning of a new session (**Extended Data Fig. 2d-f**), suggesting that transitions from goal-directed to habitual behavior may be supported by offline processing.

**Figure 2.**
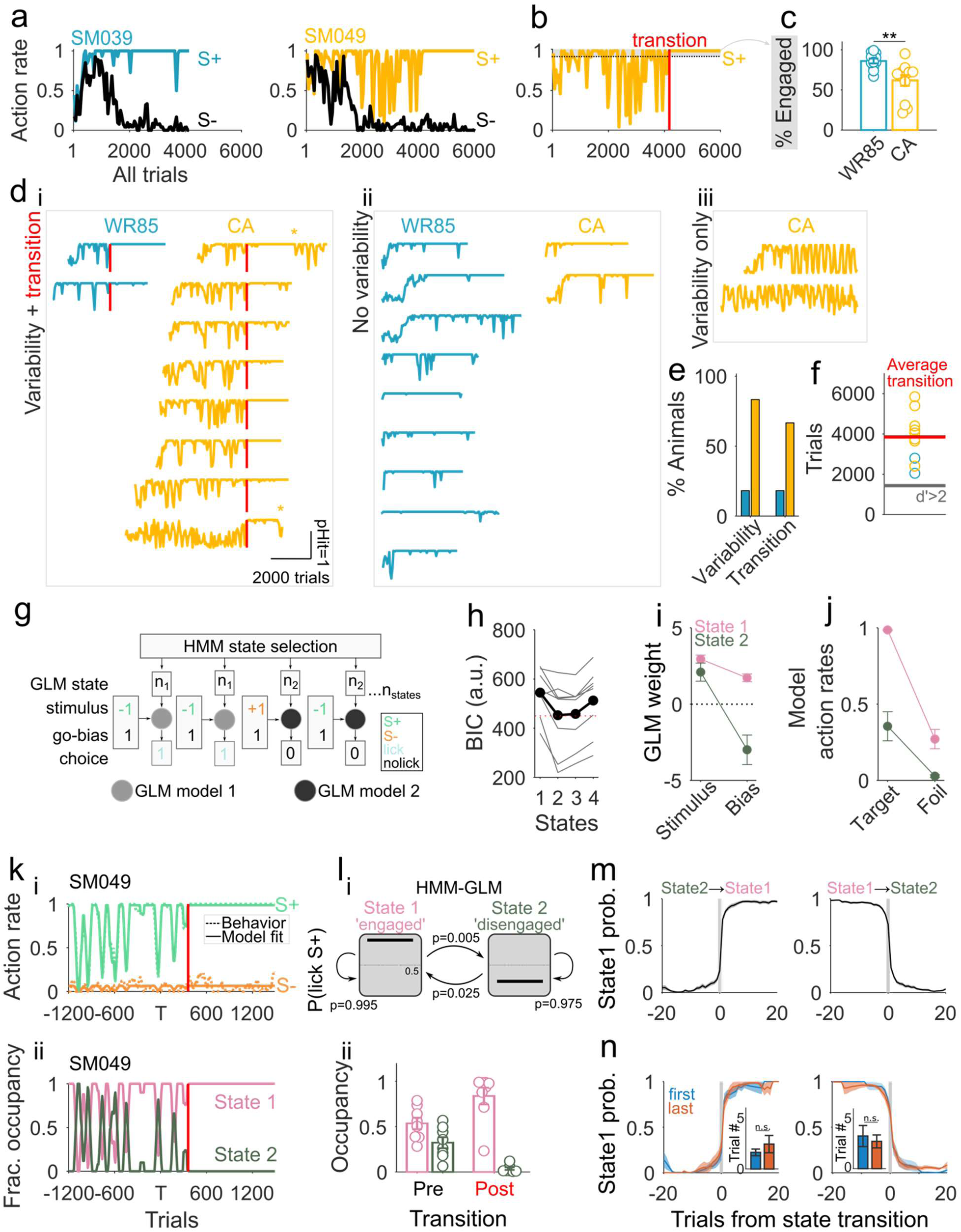
Abrupt state-like transitions from goal-directed to habitual behavior appear spontaneously within individual animals. **a,** Hit and FA action rates for example WR85 (left) and a CA (right) animals. **b,** Hit rate of CA example animal (in a) showing periods of high (gray shadow) and low engagement, followed by a spontaneous transition (red vertical line) to low hit rate variability. **c,** CA mice have significantly less engaged blocks compared to WR85 (Wilcoxon rank-sum test p=0.0028). **d_i_,** Most CA mice (8/12, 66.7%) showed hit-rate variability and the presence of a transition, while only a low percentage of WR85 mice did (2/11, 18.2%). Some CA mice that transitioned to low variability hit rate, seemed to transition to high variability after a while (asterisks). **d_ii_,** Most WR85 mice (9/11, 81.8%) showed no hit rate variability, while only a few CA mice did (2/12, 16.7%). **d_iii_,** Few CA mice showed initial hit rate variability, but never transitioned to low variability (2/12, 16.7%). **e,** Overall, most CA mice (10/12, 83.3%) showed high action rate variability, while only 18.2% of WR85 did. A high percentage of CA mice also showed transitions (66.7%), while only 18.2% of WR85 mice did (n=11 WR85 mice and n=12 CA mice). **f,** From all animals that transitioned, the average transition session is Session 13, which occurs thousands of trials after reaching expert performance (session 6) (n=11 WR85 mice and n=12 CA mice). **g,** An HMM-GLM was used to model behavioral data, which allows us to analyze the state-like nature of transitions. **h,** An HMM-GLM with two states provides the best fit for most of the CA animals based on a BIC analysis (n=8 CA mice). **i,** Both states are equally stimulus-driven, but state 2 is characterized by a NoGo, or disengaged bias (n=8 CA mice). **j,** State 1 (pink) is highly stimulus selective between target and foil trials with high engagement, while State 2 (green) shows overall task disengagement (n=8 CA mice). **k,** A CA exemplar shows that the two-state HMM-GLM model accurately recapitulates the behavior (top), and the two states govern the pre-transition phase, while only one state becomes explanatory of the post-transition phase. **l_i_,** The GLM states reflect transitions between an engaged state (state 1, pink, hit rate = 0.98) and a disengaged state (state2, olive, hit rate = 0.35). Across all mice, the HMM-GLM model predicted that the probability of staying in engaged or disengaged state (trial-by-trial) is 0.995 and 0.975 respectively, whereas the transition probability between states is 0.005 (engaged to disengaged) and 0.025 (disengaged to engaged). **l_ii_,** State occupancy pre-transition (black) is approximately divided 50%-50% between State 1 and State 2, while post-transition (red), State 1 dominates (n=8 CA mice). **m,** NoGo to Go (State 2 → State 1) transitions and Go to NoGo (State 1 → State 2) transitions happening in the goal-directed phase, are abrupt (n=8 CA mice). **n,** We observed no differences between the first (blue, belonging to the goal-directed phase), and last (red, belonging to goal-directed to habitual) transitions in abruptness. Both happen within 1-4 trials (n=8 CA mice).

Our analysis thus far required categorization of behavior based on pre-defined criteria (low versus high variability, pre- versus post-transition) and experimenter-defined parameters (**Extended Data Fig. 3a-d**). We sought to test whether a bottom-up, model-based approach could identify behavioral ‘states’ in an unbiased, and trial-by-trial, manner. To do this, we applied a generalized linear model that incorporates a hidden Markov process (HMM-GLM)^6^ on trial-by-trial behavioral data after animals reached expert discrimination performance (**Fig. 2g**). The HMM-GLM identified two states that best described the behavior in expert animals, as defined by the lowest cross-validated Bayesian Information Criterion value (BIC) (**Fig. 2h-i**). Both states were sensitive to the stimulus but with distinct action biases. State 1 (pink) exhibited high engagement, evident by a high bias and high hit rate (i.e. action rate on target trials), while State 2 (dark green) exhibited a strong disengagement, evident by a low bias and low hit rate (**Fig. 2i-j**). The HMM-GLM (solid line) accurately recapitulated the behavioral data (**Fig. 2k_i_**, and **Extended Data Fig. 3e**). Interestingly, before the transition, CA mice regularly switched between the two states both within and across sessions. After the habitual transition, however, State 1 (‘engaged’) dominated behavioral performance (**Fig. 2k_ii_-l**, and **Extended Data Fig. 3e**). We then used the HMM-GLM model to predict the transition in a bottom-up manner which we found to be similar to the behaviorally predicted one while providing greater temporal specificity (**Extended Data Fig. 3f**) (Wilcoxon rank sum test, p=0.74). The model-defined shifts from ‘Engaged’ to ‘Disengaged’ states before the habitual transition were strikingly abrupt (**Fig. 2m**). Moreover, the last transition, reflecting the putative transition between goal-directed and habitual behavior, was as abrupt as the earlier fluctuations (**Fig. 2n**, ‘last’). These state transitions occurred within less than 5 trials, and for most animals, they occurred at the very beginning of a session (**Extended Data Fig. 3g-i**) further implicating a role for offline processing in the transition of behavioral control. Thus, both quantification of behavioral data and model-based approaches using the HMM-GLM converge on the abruptness of the identified habitual transition (**Extended Data Fig. 3a-f**).

### Licking microstructures demonstrate automaticity post-transition

One alternative interpretation of these results is that animals abruptly transitioned to a higher vigilance state in which they exploit their task knowledge in a goal-directed manner, rather than a transition to habitual decision-making. In skill-based learning, motor patterns of ‘automaticity’ can be used as evidence for the formation of habits^29^. We analyzed lick microstructures in detail to determine the extent to which transitions in hit-rate variability were concomitant with changes in motor automaticity. Before transitions, CA mice exhibited highly variable lick microstructures in comparison to WR85 mice (**Fig. 3a**, top). Post-transition, however, three aspects of their licking behavior abruptly appear: a uniform lick stereotypy (**Fig. 3a**, bottom), an increase in consummatory licks (**Fig. 3b-c**, bottom), and a reduction in reaction time (**Fig. 3d**). These patterns were consistent across trials and sessions after the habitual transition demonstrating the simultaneous appearance of motor automaticity.

**Figure 3.**
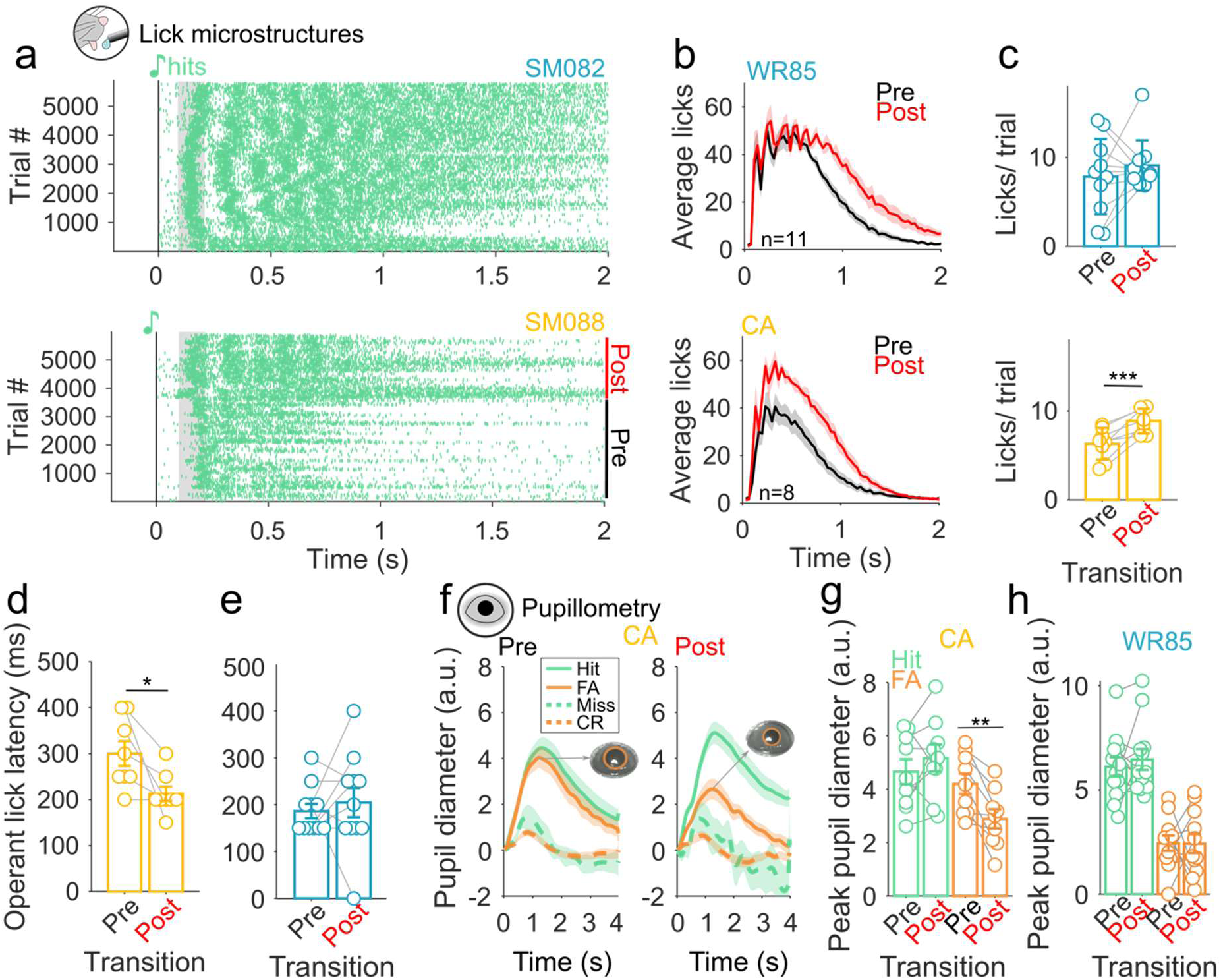
Motor automaticity and error-related pupillary signatures appear concomitant with habit transitions. **a,** Exemplar lick raster plots of a WR85 (upper) and a CA (lower) animal, showing individual licks (green lines) to target tones throughout training. Tone onset (0, black note) is followed by a dead period (100ms) and the presence of operant licks (gray rectangle). **b,** CA mice show a strong increase in post-transition number of consummatory licks (bottom) compared to WR85 mice (top). **c,** The average number of licks per trial is significantly higher in CA mice post-transition compared to pre (bottom, p=0.0019) in comparison with WR85 mice (top, paired t-test, p=0.41) (n=11 WR85 mice and n=8 CA mice). **d,** A significant reduction in the operant lick latency is observed post-transition (paired t-test, p=0.016) (n=8 CA mice). **e,** No changes in operant lick latency for WR85 mice (paired t-test, p=0.59) (n=11 WR85 mice). **f,** Evoked pupil dilation is significantly reduced for false alarms (orange) post-transition compared to evoked responses during hits (green) (paired t-test, p=0.0019) (n=9 recording days in total of n=3 CA mice for pre and post-transition). **g,** A significant difference was observed in FA evoked pupil dilation between pre and post transition in CA mice (non-parametric paired t-test, p=0.0039) but not for hits (non-parametric paired t-test, p=0.25) (n=9 CA datapoints, from 3 mice, corresponding to 3 days pre and 3 days post transition). **h,** No differences were observed in tone evoked pupil dilation during FA (non-parametric paired t-test, p=0.90), or Hits (non-parametric paired t-test, p=0.99) between pre and post transition in WR85 mice (n=12 WR85 4 mice, corresponding to 3 days pre and 3 days post transition).

### The underlying decision process is reflected in pupillary dynamics

With the ability to pinpoint the transition from goal-directed to habitual behavior, we next sought to examine whether the decision controller being used could be inferred from changes in pupillary dynamics. When behavior is under goal-directed control, the execution of an action is driven by the expectation of reward (the action-outcome contingency). In contrast, when behavior is under habitual control, the execution of the action is driven by the presence of a cue (the stimulus-action contingency) and errors are likely due to ‘slips of action’ with little relation to the outcome expectation^30^. One tool commonly used as a biomarker of decision-making processes is the pupillary response, as pupil dynamics reflect choice^31^ and track value-based decision-making^32^. Changes in pupil size are associated with increased reward magnitude, task investment or effort^33,34^, and reward expectation^35^. Here, we hypothesized that habitual behavior should recruit less cognitive effort and lower sensitivity to errors and would be reflected in changes in pupillary dynamics.

To test this, we recorded and quantified trial-level pupil responses in a subset of CA (n=3) and WR85 (n=4) mice. We aligned our phasic, task-evoked pupil measurements to the transition trial (empirically calculated and confirmed with the HMM-GLM) in CA mice. We observed that consistent with previous work^31,36^, false alarms elicited strong pupil dilations (expert and pre-transition, **Fig. 3f**, left and **Fig. 3g**, left). During habitual behavior (post-transition), however, false alarms elicited a much weaker pupil dilation (**Fig. 3f**, right and **Fig. 3g**, left). These changes could not be explained by commonly reported movement-evoked pupillary response^37–40^ as the rate of false alarms (**Extended Data Fig. 4a**) and licks per false alarm (**Extended Data Fig. 4b**) were similar pre- and post-transition. This effect was not observed in WR85 mice as the underlying motivational state likely overwhelms subtle changes in pupillary dynamics (**Fig. 3h**, and **Extended Data Fig. 4c**). These data demonstrate that the differences in tone-evoked pupil dilation reflect decreased error sensitivity during habitual behavior, suggesting that behavioral control has shifted to a less deliberative and less cognitively demanding controller.

### Reward devaluation confirms timing of habit transitions

A current gold standard for assessing whether a behavior is under goal-directed or habitual control is the use of discrete, outcome devaluation sessions^28^. To test whether our volitional engagement paradigm is consistent with this method, we reasoned that all animals are likely to transition to habitual behavior but that the transition in WR85 mice is masked by their continuously high hit rates due to ceiling levels of motivation. We inferred the transition in WR85 mice using our median transition session from CA mice (session 13). We used discrete satiety test sessions before and after this inferred transition. In these sessions, mice had *ad libitum* access to plain water for 10 minutes (**Fig. 4a_i_**) prior to performing the go/no-go task (60 trials). Under goal-directed control, behavior is expected to be sensitive to satiety (**Fig. 4a_ii_**, left) while under habitual control, behavior is thought to be insensitive to satiety (**Fig. 4a_ii_**, right). We found that pre-transition (10^th^ session), WR85 mice were highly sensitive to reward devaluation, abolishing responses (**Fig. 4b**) (Session 10, non-devalued vs devalued paired t-test, p=0.0000019), while post-transition, these same animals were less sensitive (**Fig. 4b**, devalued session 10 vs devalued session 15, paired t-test p=0.0072). Importantly, we observed no differences in non-devalued (ND) actions rates between pre- and post-transition (**Fig. 4b**, ND session 10 vs ND session 15 paired t-test p=0.25). This effect was independent of water consumption during the devaluation sessions (**Extended Data Fig. 5a**). To further validate these results obtained in WR85 mice, we tested within-session satiety in CA mice (satiety-based devaluation was not possible in CA mice, as they only intermittently drank plain water even when it was freely available, similar to lick training, **Fig. 1**). During goal-directed behavior (session 10), CA mice exhibited reductions in hit rate when comparing the beginning to the end of the session (**Fig. 4c**, black) (paired t-test p=0.038), suggesting a session-level impact of satiety. Interestingly, during habitual behavior (session 15), we observed no such reduction in hit rate (**Fig. 4c**, red) (paired t-test, p=0.24), suggesting no impact of session-level satiety. These data help to validate the volitional engagement paradigm as a means to assess the underlying decision process.

**Figure 4.**
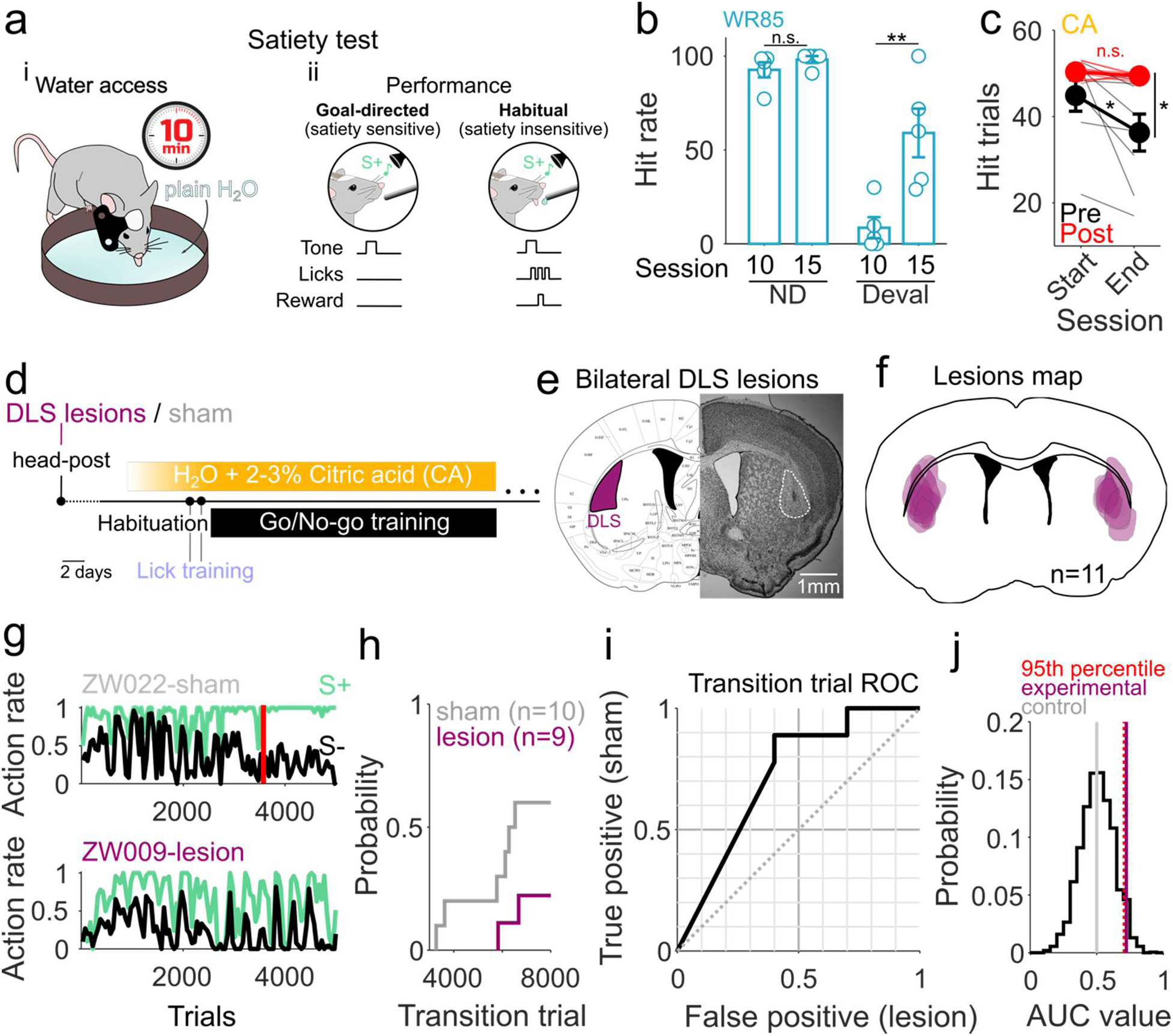
Bilateral DLS lesions delay or block transition to habitual behavior. **a_i_**, Satiety test. WR85 mice receive free access to plain water for 10 minutes right before training. **a_ii_,** During training, if goal-directed, mice are expected to reduce their action rates since they are sensitive to satiety. In habitual mode, mice tend to maintain a high action rate since they have developed a habit and become less or insensitive to satiety. **b,** Hit rates during Non-devalued (ND, sessions 9) and devalued (Deval, session 10) trials for WR85 mice. Pre-transition (session 10) mice show high sensitivity to devaluation (ND vs Deval Session 10, paired t-test, p=0.0000019). Hit rates during Non-devalued (ND, session 14) and devalued (Deval, session 15) trials for WR85 mice. Post-transition (session 15) mice show reduced sensitivity to devaluation (ND vs Deval Session 15, paired t-test, p=0.017), with action rates significantly different from the pre-transition phase (paired t-test, **p=0.0072) (n=5 WR85 mice). **c,** Intra-session satiety is observed in CA mice pre-transition but not post-transition. While animals pre transition significantly reduce their licking to S+ by the end of a session (indicating they have been sated), post transition, these same animals show no differences. The number of target trials with licks is significantly different between the start and end of a session pre-transition (black, paired t-test, p=0.038), but not post-transition (red, paired t-test, p=0.24). A significant difference is also observed between pre- and post-transition hit trials at the end of a session (end, paired t-test, p=0.017), while there are no differences at the start of a session (start, paired t-test, p=0.15) (n=8 CA mice). **d,** Lesion protocol. Mice undergo an NMDA lesion or sham right before head-post implantation. The rest of the water restriction, habituation and training protocol is as shown in Figure 1a. **e,** Lesion exemplar. Coronal view of the bilateral lesion sites at the DLS. **f,** Lesion map of all animals (n=11 mice), showing homogeneous and overlapping lesions. **g,** Action rate exemplars for sham (top, gray) and lesioned (bottom, purple) animals. **h,** Cumulative distribution of transition trial for animals that showed behavioral variability (n=10 sham and n=9 lesioned mice). While 60% of sham animals show a transition, only 20% of lesioned animals do. **i,** ROC curve built from transitions probabilities in g. **j,** AUC values for shuffled labels in our experimental groups. The difference in AUC between sham and lesioned mice (purple line) falls into the 95^th^ significance percentile (red dotted line), compared to the difference between control animals (gray line).

### Bilateral lesions to the DLS prevent transitions to habitual behavior

The two decision processes (goal-directed and habitual) are thought to be sub-served by distinct neural circuits in the dorsal striatum^41^. The dorsomedial striatum (DMS) and dorsolateral striatum (DLS) enable goal-directed and habitual behavior, respectively^42,43^. Using standard outcome devaluation procedures, rodents with lesions to the DLS persist in goal-directed mode (i.e. when they should be sensitive to outcome devaluation) even after significant amounts of overtraining^7^. Thus, DLS lesions provide a powerful and orthogonal approach to test the validity of our volitional engagement paradigm in assessing the underlying decision mode. To do this, we bilaterally lesioned the DLS in CA mice (NMDA 20µg/µl, 100nl/site) before head post implantation and behavioral training (**Fig 4c**). All DLS-lesioned CA mice had visible, localized, and overlapping lesions (**Fig. 4d-e**, and **Extended Data Fig. 5b**), while shams did not (**Extended Data Fig. 5c-d**). DLS-lesioned CA mice exhibited high variability in hit rates that persisted for much longer than in sham CA mice (**Fig 4f**, and **Extended Data Fig. 5e**) and lesioned animals rarely transitioned to habitual behavior (**Fig. 4g-i**, and **Extended Data Fig. 5e**) (60% sham vs 20% lesioned). Importantly, CA mice with DLS lesions exhibited no significant deficits in the learning of the task contingencies (**Extended Data Fig. 5f-g**). These data offer independent evidence, through the manipulation of habit-relevant striatal circuits, that the transitions we observe are indeed genuine transitions from goal-directed to habitual behavior.

## Discussion

A fundamental tenet of animal behavior is the existence of multiple ‘controllers’ that govern decision-making. One prevailing framework is that instrumental decisions come about from two distinct processes: goal-directed and habitual^44^. In rodent-based learning paradigms, goal-directed behaviors are thought to become habitual upon overtraining^45–47^. The goal-directed system dominates early in learning when animals will ‘work’ for food or water because the outcome itself is desirable. This requires both a representation of the action-outcome contingency and the recognition of the outcome as a motivational incentive. When under goal-directed control, behavioral decisions are ‘value-based’ and flexible but also cognitively demanding^48^. The habitual system is thought to take over during overtraining to simplify the decision process and reduce cognitive complexity by shifting to a stimulus-response mode of behavior^48^. When under habitual control, behavioral decisions can be considered ‘value-less’ and inflexible but also less cognitively demanding^49^. Over the past 50 years, behavioral, neural, and theoretical support for these two distinct decision processes (but also the complexity of their interaction) has grown largely due to behavioral manipulations, including outcome devaluation, contingency degradation, and the use of different reinforcement schedules. These behavioral tools have been invaluable to gain a deeper understanding of the behavioral, neural, and theoretical basis of the multiple systems controlling decision-making. Nevertheless, the extent to which discrete measures of sensitivity to outcome devaluation sufficiently distinguishes goal-directed from habitual control is still under scrutiny^50,51^ as sensitivity to outcome devaluation can also be triggered by unexpected cues^52^ and in situations where habits are expected to form^53^. More broadly, the current methodologies remain fundamentally limited in their temporal resolution and individual specificity, limiting the assessment of nature, timing, and properties of habit formation^19,50^ in individual animals. As a result, an essential question first posed by Clark Hull in 1943 has remained unresolved: ‘Is this transition abrupt, or is it gradual and progressive?’

Here, we lay out an approach that allows real-time assessment of the underlying decision process without the need for discrete testing sessions and without the implementation of specified training schedules to bias decision modes^26,54,55^. Rather than binarize motivation into motivated (non-devalued) and un-motivated (devalued), we sought to mimic naturalistic motivation levels. We hypothesized that by shifting an animals’ desire for an outcome from a *need* (survival) to a *preference* (palatability), animals would gain agency on their motivation to engage in a task based on the desire for an outcome (in our case, plain water droplets). Here, we show one approach to such a ‘volitional engagement’ paradigm by giving animals *ad libitum* access to CA water^4,5^ in the home cage, which reduces the motivation for plain water (**Fig. 1b-d**). In goal-directed mode, animals volitionally engage and disengage from the task, reflected in the behavior as state-like switching early in learning (see **Fig. 2a** and **Fig. 2d**). As behavior becomes habitual, animals shift to constant engagement, behaviorally observed as an abrupt shift to a constantly high action rate (**Fig. 2a**, **Fig. 2d**, **Fig. 2m**, and **Extended Data Fig. 3a-e**), contradicting the assumption that habit expression is gradual. These transitions occurred thousands of trials after reaching expert discrimination performance (**Fig. 2f**) but at different time points for individual animals.

We then used orthogonal measurements of motor automaticity, pupillary dynamics, sensitivity to traditional outcome devaluation, and lesions of the DLS to confirm that our observations reflected a transition to habitual decision-making versus differences in vigilance or discrimination ability. In other words, behavioral automaticity and reduced error sensitivity occurs concomitant with habit formation in our paradigm. While automaticity alone might be a reductionist perspective in habit formation^53^, the composite picture across behavioral and neural approaches in our study points toward the habitual nature of behavior post-transition in volitionally engaged mice. This novel approach— which we term ‘volitional engagement’—adds a powerful *en passant* tool to study habit formation and perseverance.

Pinpointing the precise nature of this transition under naturalistic motivational conditions provides a powerful tool to explore the psychological and neural basis of habit formation in real-time. An abrupt ‘insight-like’ transition suggests that a higher-level process operates in conjunction with lower-level associative processes. Our findings challenge the notion that habit expression is cumulative, with its likelihood increasing incrementally with each successive reinforcement. We demonstrate that habits appear nearly instantaneously (within 5 trials) with animals having received vastly different levels of reinforcement during training. This suggests that a separate higher-level process arbitrates between goal-directed and habitual control. Factors such as cognitive demand or environmental uncertainty likely contribute to when the commitment emerges to solve the task in a simple and inflexible manner, activating an otherwise dormant habitual controller. The habit becomes instantiated precisely when animals choose to use the habitual controller, whether they explicitly want the water or not. In this view, animals under habitual control can still internally ‘experience’ periods of goal-directedness (i.e., they still want plain water sometimes). In addition, the higher-level nature of the choice suggests that habitual control need not be permanent. Interestingly, some animals reverted to goal-directed mode after several sessions in habit mode (see **Fig. 2d**, yellow asterisks and **Extended Data Fig. 5e**, asterisks), suggesting that transitions to habitual decision-making are not intrinsically permanent. This higher-level process that controls the switch may also help explain why techniques such as outcome devaluation yield conflicting results in rodents^56^ (due to individual variability in when transitions occur) and remain largely ineffective in humans^19^ (given a more complex interaction between cognitive and motivational drivers).

The current consensus, though contentious^57^, is that habitual and goal-directed behavior are supported by the DLS and DMS, respectively^42^, but studies utilizing discrete satiety test sessions or experimenter-defined overtraining periods yield confounding and even conflicting results: some evidence argues that DLS activity changes rapidly before the behavior onset of a habit^58^, with others finding that the change is more gradual and closely aligned with a behavioral change^59^. Recent reports even observe an eroding distinction in the control of actions between the DLS and DMS as training progressed^57^. The spontaneous and abrupt appearance of habitual control provides a behavioral marker upon which to identify ‘switch-like’ activity in the underlying neural circuits^19^. The possibility of a higher-level process that arbitrates between goal-directed and habitual control points to regions such as premotor or prefrontal cortical areas^17^, which have bidirectional interactions with the DMS and DLS. Alternatively, this arbitration process may be governed in the striatum itself. Identifying the neural circuit dynamics that govern this transition remains an important area for future investigation.

Finally, our discovery of abrupt transitions to habitual behavior may inform distinct interventional strategies in habit disorders in humans. Rather than relying on gradual exposure and/or systematic desensitization, there might be value in interventions that combine such associative strategies with cognitive control strategies. Our data suggest that it may be possible to predict when transitions will occur and if such predictions are possible and can be extended from rodents to humans, it could provide a powerful tool to interfere or manipulate the emergence of maladaptive habits.

## Methods

### Animals

All mice were housed in standard plastic cages with 1-4 littermates and kept in a 12-h/12-h light/dark cycle (10:30 am / 10:30 pm) with controlled temperature (19.5-22°C) and humidity (35-38%). All the mice used in this study were male C57BL/6J from Jackson Labs (strain 664) with an age of 11.61±0.21 (average ± SEM) weeks at the start of training). All the experimental and surgical procedures were approved and performed in accordance with the Johns Hopkins University IACUC protocol (license # MO20A272).

### Surgical procedures

Mice were anesthetized with isoflurane (5.0% at induction, 1.5-2.5% during surgery) and placed on a stereotactic apparatus (Kopf). The hair over their skull was removed with hair removal cream and the area disinfected with betadine. The skin over the skull was removed and the area was cleaned of connective tissue with 3% H_2_O_2_. A custom-made stainless-steel head-post was fixed onto the exposed skull with C&B Metabond dental cement (Parkell). The animals were given 1-3 days to recover. Mice that underwent bilateral DLS excitotoxic lesions or sham injections, received NMDA (Sigma Aldrich, 20µg/µl NMDA in PBS1x with 10% glycerol) or vehicle (PBS1x with 10% glycerol) respectively (with a Hamilton syring and a Havard Apparatus Pump 11 Elite, 100nL/injection-site at a 70nL/min). The injections were made at AP+1.0mm, DV±2.6mm, ML-2.8mm via burr holes which were sealed with Jet Denture Repair Acrylic (Lang Dental) prior to headpost implantation.

### Histology

At the end of the DLS lesion experiment, the brains of all animals were obtained via transcardiac perfusion^60^ and stored in 4% paraformaldehyde solution in PBS1x overnight. After further dehydration in 30% sucrose (Sigma-Aldrich), the brains were frozen in OCT gel (Tissue-Tek®) and sliced using a cryostat (Leica) into 50um slices. The slices were mounted on gelatin-coated slides (FD Neuro) and left in room temperature to dry overnight. The following day, the slides were stained using 1% cresyl violet (Sigma-Aldrich) solution and cover glasses were placed and fixated with Permount mounting medium (Fisher Chemical). The slides were imaged under Brightfield settings in a Zeiss upright microscope (Axio Zoom.V16).

### Habituation and water restriction paradigms

After recovery from surgery, animals were handled and habituated prior to the start of training for at least 10 days based on previous studies^61^. Head-fixed experimental CA mice and their littermate controls (WR85) underwent the same surgical, habituation and testing procedure. Animals were handled by the experimenter/s at increasing times every day, exposed, and habituated to the head fixation station. The different water restriction paradigms started after at least 3 days of handling. The standard water restriction (WR85) protocol prevented the animals from accessing water in their home cage. The mice were weighed daily, and a limited amount of water (∼1.0 mL) was given individually to maintain 80-85% of their original weight. For naturalistic water restriction with citric acid (CA), animals had *ad libitum* access in their home cage to a bottle of tap water with citric acid dissolved. The mice were slowly introduced to the taste of CA, increasing its concentration daily from 0.5% CA to 1-3%, and adjusted accordingly within this range to keep the animals at ∼95% of their original weights.

### Behavioral training

All behavioral training was done using Bpod State Machines (r1 or r2, Sanworks). After habituation, mice underwent an initial instrumental training phase where they were head-fixed and trained to lick from a lick tube by rewarding each lick with a drop of water (3 µl). There was no tone stimulus presented during lick training. The lick training session ended either when 1 ml of water was consumed, or session had reached 30 minutes. On a subsequent session, mice began training on a go/no-go auditory task. Behavioral events (trial structure, stimulus and reward delivery, lick detection) were controlled and stored using a custom-written MATLAB program (2018b, The MathWorks) interfacing with the Bpod, an electrostatic speaker driver (E1, TDT) and an infrared beam for lick detection. In a subset of animals, facial movements and pupil size were measured with a Raspberry Pi (3B) and a Raspberry Pi camera module (NoIR v2) coupled with a Bright-Pi infrared LED array (PiSupply). Mice were head-fixed inside a Plexiglass tube facing a lick-tube. A free field electrostatic speaker (ES1, TDT) was located ∼5 cm from the animal’s left ear and each sound (either 4757 Hz or 8000 Hz, as target or foil stimuli) was calibrated to an intensity of 60-62 dB (SPL). The pupil camera and IR LED array were positioned ∼6 cm away from the animal’s face in a 60-degree angle. Everything was enclosed in a custom-made sound-attenuated box. Target and foil tones were pseudo-randomly ordered (equilibrated every 20 trials). Each trial consisted of a pre-stimulus no-lick period (2 s), stimulus presentation (100 ms), delay (100 ms), response period (2 s) and variable inter-trial interval (ITI). Typically, mice were trained for ∼300-320 trials per with a short block of 20 non-reinforced trials interleaved in the middle of the session^62,63^. Training lasted for a maximum of 30 days.

### Behavioral analysis

Individual-animal action rates were measured in blocks of 50 trials to obtain hit and false-alarm rates in discrete but small blocks that allowed us to observe behavioral variability. Behavioral discriminability was calculated using the z-scored hit rate minus the z-scored false-alarm rate (d’). To avoid infinite values when rates of 1 or 0 are present, the values were corrected by 1-1/2N or 1/2N respectively, where N corresponds to the number of trials. For all the data presented in this paper, we considered animals’ to have effectively learned the task by calculating d’ during non-reinforced trial blocks, which has previously been demonstrated as an accurate measure of task acquisition^62^. Only mice with a d’ > 2 during these non-reinforced trial blocks were included in the analysis (corresponding to 45 out of 48 mice tested, 1 WR85, 1 CA-sham and 1 CA-lesioned mice did not learn the task and thus were excluded).

### HMM-GLM model implementation

We fit a GLM-HMM model to trial-by-trial choices of each mouse from 4 days before putative habitual transitions to 4 days after the putative transition (9 days in total). Each state in HMM contains a Bernoulli GLM defined by a weight vector specifying how stimulus inputs and bias are integrated in that state. The model was fit using a previously published expectation-maximization (EM) algorithm^6^. To identify the optimal number of states, we evaluated the cross-validated BIC by fitting choice data from the 5th day before and after habitual transition. A 2-state model was sufficient to explain the choice behavior of six animals, capturing an engaged state and a disengaged state, whereas a 3-state model was needed for two animals, capturing an additional low-discrimination state. For these two animals, we focused only on the engaged and the disengaged state in subsequent analysis. To compute state occupancy, we first inferred the behaviorally dominant state as the state with the highest probability in each trial, and then calculated the percentage of trials that a state is dominant in a 50-trial bin. We inferred the habitual transition by identifying the last trial bin where the occupancy of disengaged state was above a threshold of 30%. The number of trials needed for transitions between engaged and disengaged state is calculated by the number of trials needed for the dominant state to reach 75% probability after the transition. To compare the model-inferred transitions with behavioral data, we quantified the slope of inferred state probability by the GLM (z-scored) at the trial of state transition, compared to the slope of hit rate changes during state transitions (z-scored), quantified using various bin sizes around the transition.

### Preprocessing of pupillometry data

20 minutes long pupillometry videos (n=5) were taken as the training dataset for a DeepLabCut^64,65^ (DLC) pre-trained model (resnet_50). Manual labeling of pupil contour consisted of 8 points (up, down, left, right, up-left, up-right, down-left, down-right) across 180 randomly selected frames. The network was trained for 564,000 iterations until the loss rate plateaued. The final network was used to analyze the pupillometry videos from the experiment. Custom MATLAB code (The MathWorks, 2019b or 2022a) was then used to remove blink artifact, reconstruct pupil diameter, and apply a low pass filter (3 Hz) to the data. Individual trials for individual animals were normalized to the median pupil diameter during correct reject trials per session.

### Statistical Analysis

All analyses were performed using custom-written MATLAB code (The MathWorks, 2019b or 2022a). All datasets were tested for normality using a one-sample Kolmogorov-Smirnov test; then, parametric or non-parametric statistical tests were applied accordingly. Two-sample t-tests were used for parametric data, and Wilcoxon rank sum tests were used for non-parametric data. Where required, paired comparisons were made. For multiple comparison analyses, 2-way ANOVA was performed. To build a Receiver Operating Characteristic (ROC curve) (**Fig. 4h**) we used the transition probability of lesioned and sham animals to obtain the area under the curve (AUC) and generate a shuffled probability distribution to statistically test our experimental animals’ distribution difference. Significance was determined as the difference in AUC value between lesioned and sham animals when it fell beyond the 95^th^ percentile confidence interval (**Fig. 4i**). All confidence intervals correspond to α=0.05. Significance is represented as n.s. p>0.05, * p≤0.05, ** p≤0.01, *** p≤0.001, and **** p≤0.0001.

### Data reporting

Sample sizes were determined based on standard cohort sizes from relevant literature. Mouse allocation to specific groups was randomized but the experimenters were not blinded to group types.

### Reporting Summary

Further information on research design will be available in the Nature Portfolio Reporting Summary linked to this article.

## Data availability

Data will be made available upon acceptance of this manuscript.

## Code availability

No specialized software was developed for this work.

## Author information

### Contributions

SM and KVK designed the project. ZW, SM and AL performed the experiments. SM, ZZ, and RS analyzed the data. ZZ, RS, and AC performed computational modeling. SM performed final analysis, figures, and data curation. SM, ZW, and KK wrote the manuscript. KVK provided funding and supervised the project. All authors participated in results interpretation and manuscript editing.

## Acknowledgements

We thank P. Janak, A. Haith, J. Krakauer, N. Kothari, P. Holland, and Y. Cheng for helpful comments on the manuscript. We thank E. Barker and D. Udzinski for animal care taking. This work was supported by grants from the NIH R01DC018650 and R00DC015014 to KVK.

## Extended data and figures

Extended figures 1 to 5 and legends are provided.

---

**Extended Data Figure 1.**
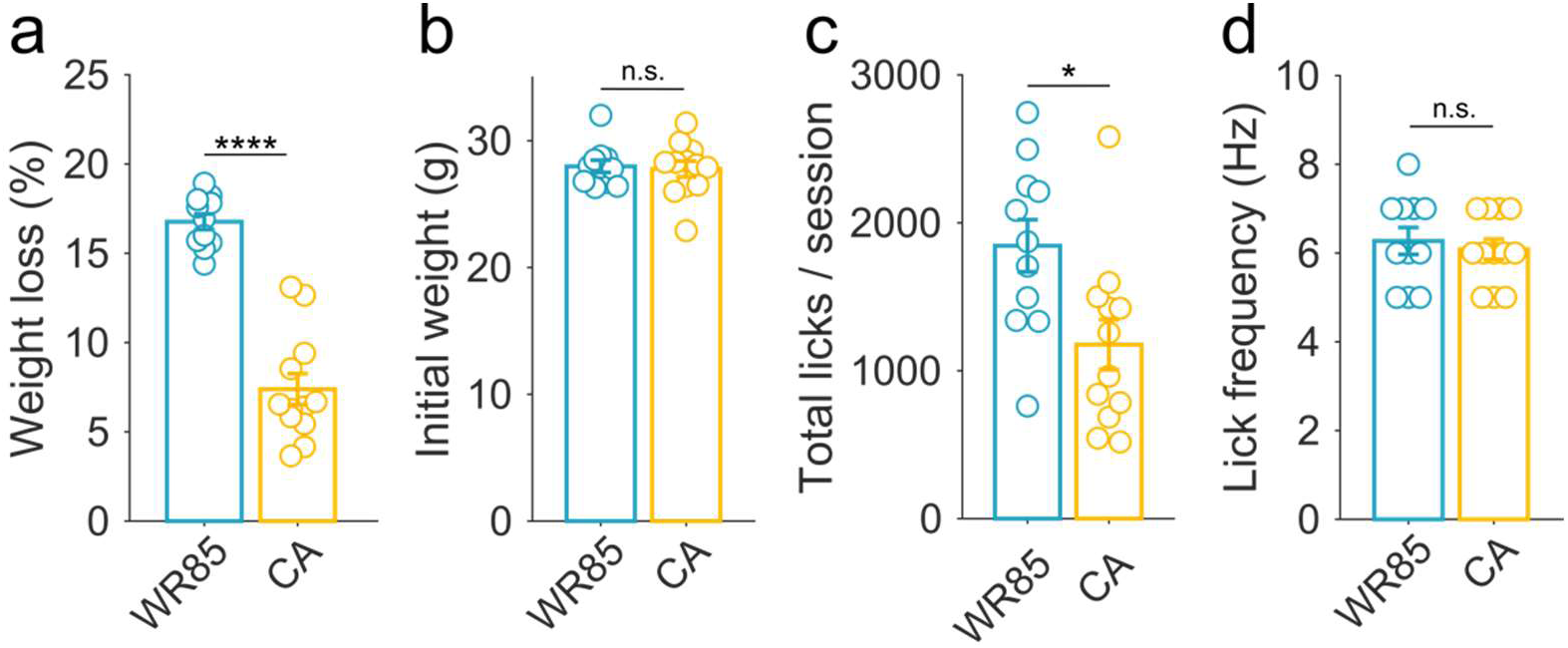
Palatability-based motivation is stable throughout training and reduces an animals’ motivation to obtain water rewards. **a,** Weight loss during go/no-go training was significantly reduced in CA mi*ce* (Wilcoxon ranksum test, p=0.000055). **b,** Animals’ original weights were not different between groups (Wilcoxon ranksum test, p=0.97). **c,** CA mice also do significantly lower number of total licks in a lick-training session (Wilcoxon rank-sum test, p=0.021). **d,** Lick vigor (frequency) was not different between groups (p=0.67). **a-d,** n=11 WR85 mice and n=12 CA mice.

**Extended Data Figure 2.**
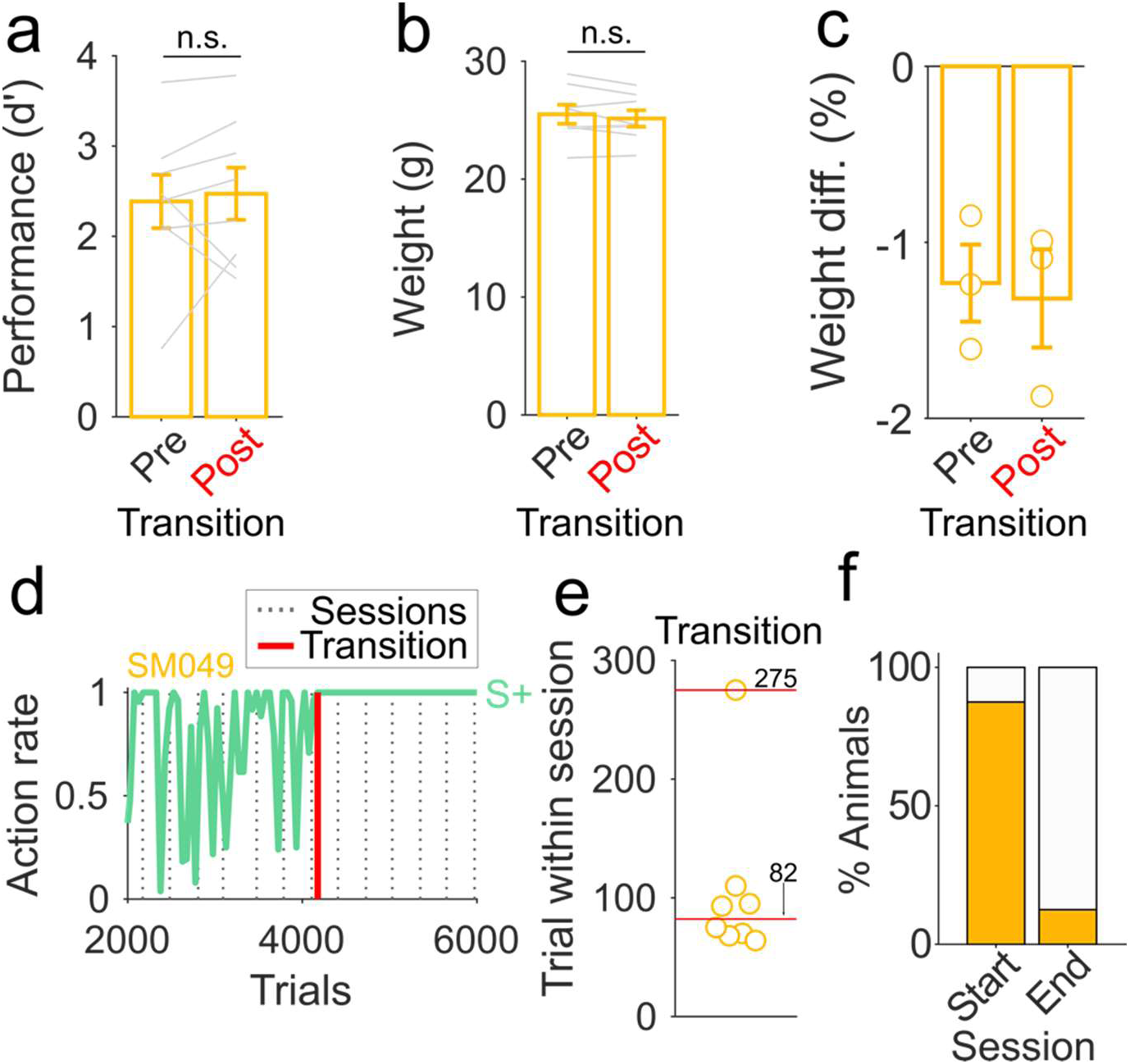
Spontaneous transitions from goal-directed to habitual behavior are independent of performance or metabolic state and occur at the beginning of a new session. **a,** No significant differences are observed between pre and pos-transition performance (Wilcoxon signed rank test, p=0.54, n=8 mice). **b,** Weights of CA mice with transitions are stable around the transition session (average of 3 sessions pre and post transition). No significant differences are observed between pre-transition weight and post transition weight (Wilcoxon signed rank test, p=0.29, n=8 mice). **c,** Weight differences between the end of a session and before the start of the next day session confirm that animals maintain similar consumption rates in their home-cage comparing pre and post-transition (Wilcoxon signed rank test, p=0.82, n=3 mice). **d,** Exemplar animal of hit rates (green) with the overlying sessions (vertical dotted gray lines) showing that the transition to habit happened close to the start of a new session. **e,** Of the 8 CA mice with transitions, 7 transitioned early in the session. These data use a 50-trial binning procedure that limits the temporal resolution, such that animals transitioning at Trial ∼80, are likely transitioning much closer to the beginning of the session. See Extended Data Figure 3g-h for a model-based estimation using trial-level data. **f,** The vast proportion of CA mice transition at the start of a new session (trials 1-150), compared to in the end of a session (trial 151-300) (n=8 mice).

**Extended Data Figure 3.**
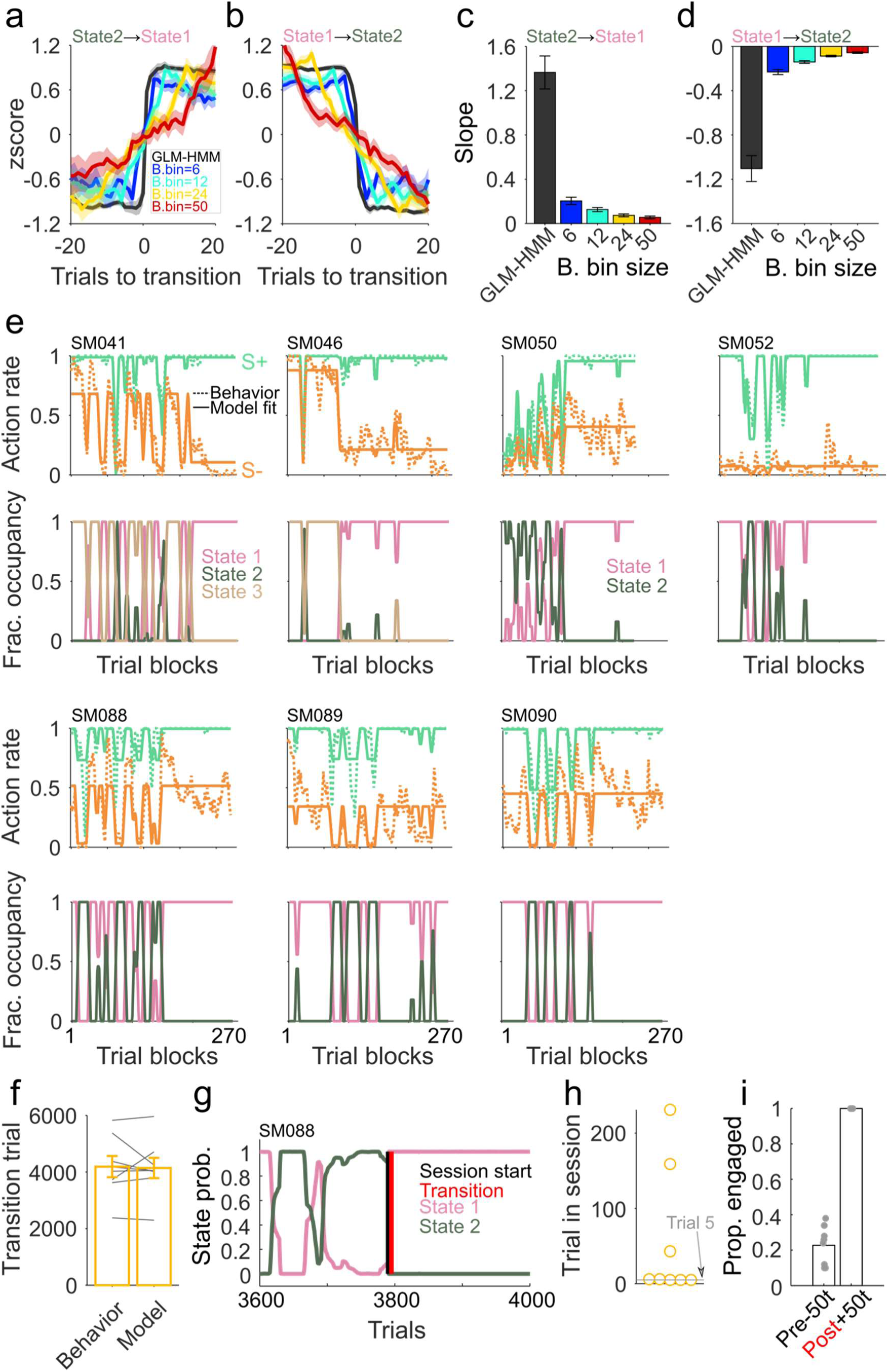
State-like transitions are observed in individual animals using an HMM-GLM model. Z-scored model state probability (black) or behavioral hit rate using bin size from 6-50 trials (blue – red) for **a,** disengaged-to-engaged and **b,** engaged-to-disengaged fluctuations. Quantification of abruptness using slope of the trajectories near the transition point for **c,** disengaged-to-engaged and **d,** engaged-to-disengaged fluctuations. **e,** Model fit plots and fraction occupancy plots for all additional CA individual animals that transitioned (n=7). **f,** The trial-by-trial nature of the model allowed us to find exact transition points, which are similar to the behaviorally identified ones using only hit rates (Wilcoxon signed rank test p=0.95). **g,** State probability for State 1 (pink), State 2 (green) for an example animal right around the transition session (black vertical line), depicting that transitions (red vertical line) typically happen right at the start of a new session. **h,** Most animals (5/8) transition within 5 trials, but others transition later in a session. **i,** 50 trials before the transition point (Pre -50t) there is very low probability of task engagement, while 50 trials after the transition (Post +50t), engagement is at its maximum.

**Extended Data Figure 4.**
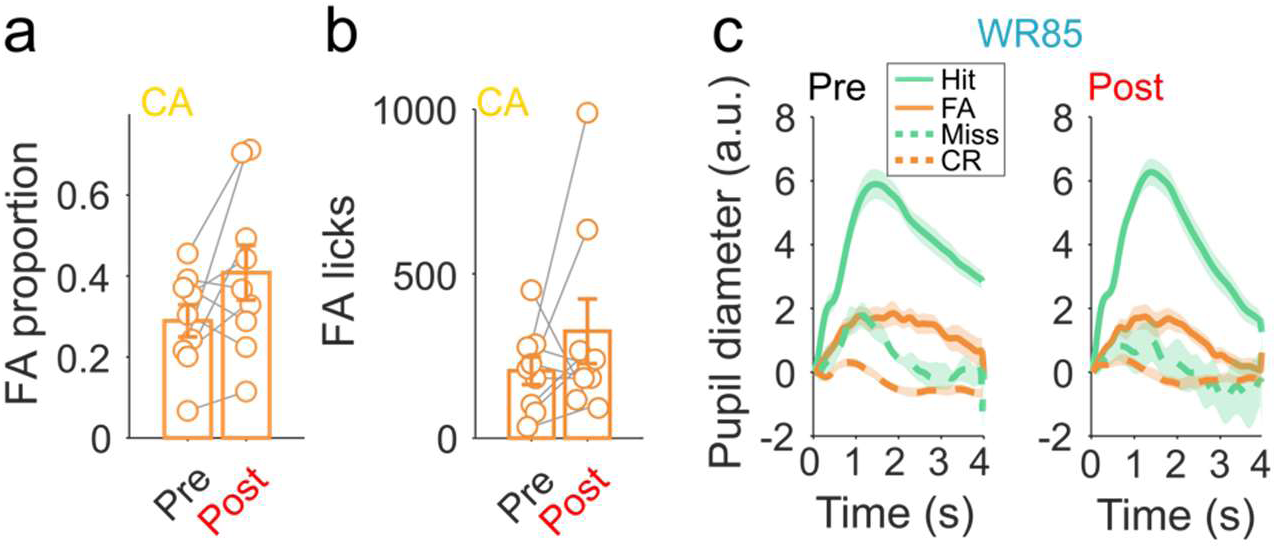
Changes in pupillary error signal between pre and post transition are only evident in CA mice and independent of movement-evoked pupillary changes. **a,** The proportion of FA in CA mice is not different pre and post-transition (Wilcoxon signed rank test, p=0.098) (n=3 days pre and 3 days post from 4 mice). **b,** No changes in the number of licks to FA were seen in CA mice pre and post-transition (Wilcoxon signed rank test, p=0.25) (n=3 days pre and 3 days post from 3 mice). **c,** Tone-evoked pupil dilation during hits or FA is not changed between pre and post-transition in WR85 mice (n=3 days pre and 3 days post from 4 mice).

**Extended Data Figure 5.**
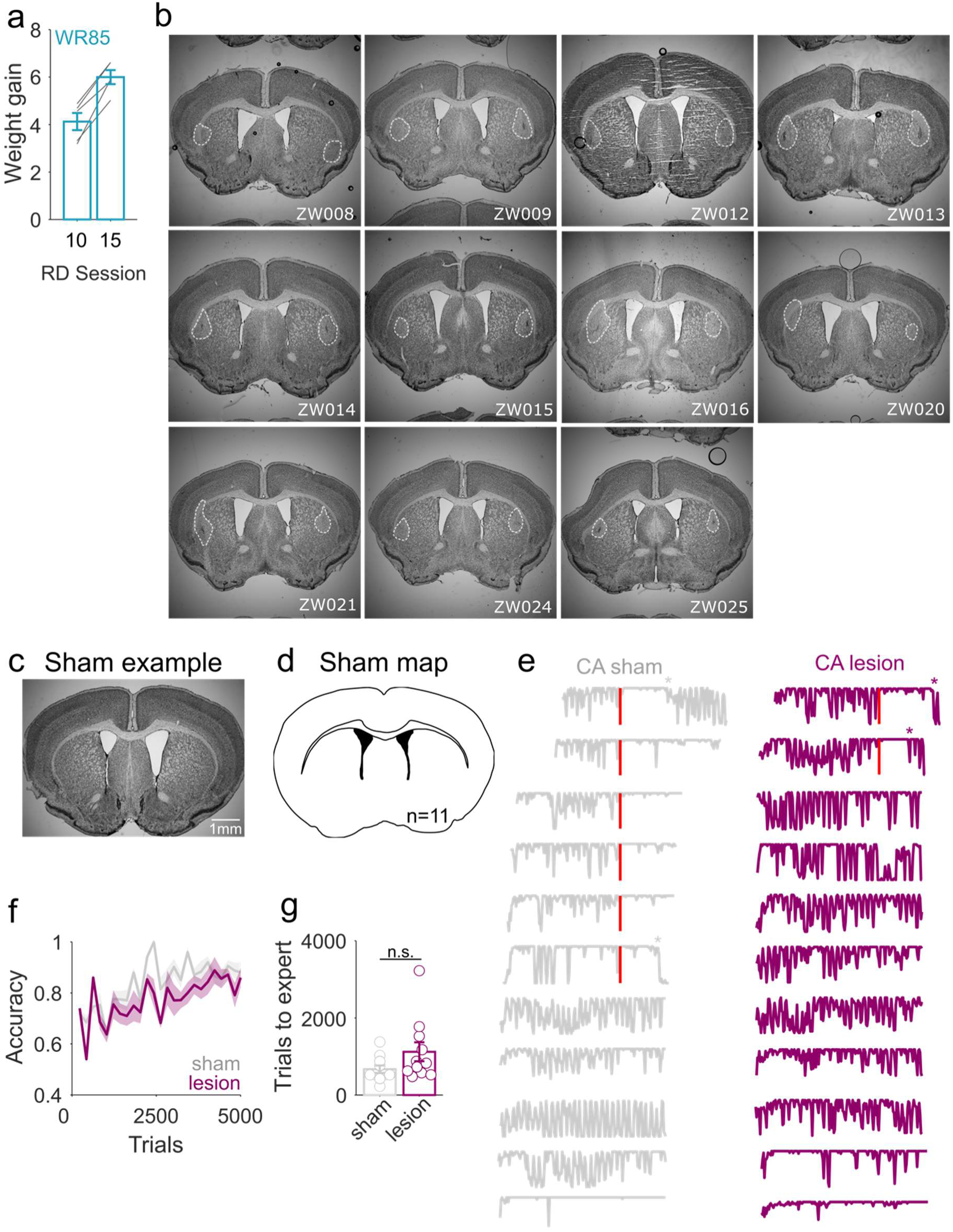
DLS lesioned animals are less likely to transition to habitual behavior. **a,** No decrease in weight between reward devaluation (RD) session 10 and 15, suggesting that differences in the increased hit rates seen in session 15 are not due to reduced water consumption during the satiety test (Wilcoxon signed rank test, p=0.0039) (n=5 WR85 mice). **b,** CA-sham exemplar showing no DLS lesions when injected bilaterally with vehicle. **c,** Map of n=11 CA-sham mice showing no lesion-like areas for any individual. **d,** All individual lesioned mice with outlined lesion area (dotted white line). **e,** Sham (gray) and lesioned (purple) individual mouse hit rates, depicting a transition to habitual behavior (red). Animals that transition back to goal-directed control are shown with an asterisk. **f,** No differences in task accuracy between sham (gray) and lesioned (purple) animals (2-way ANOVA, F(1,25)=0.43, p=0.51, interaction group x trials F(1,25)=0.5, p=0.98) (n=11 sham and n=11 lesioned mice). **g,** Similar number of trials to expert performance between sham (gray) and lesioned (purple) mice (Wilcoxon ranksum test p=0.098) (n=11 sham and n=11 lesioned mice).

